# Protein Carrier AAV

**DOI:** 10.1101/2024.08.14.607995

**Authors:** Mareike D. Hoffmann, Ryan J. Sorensen, Ajay Extross, Yungui He, Daniel Schmidt

**Affiliations:** Department of Genetics, Cell Biology & Development, University of Minnesota, Minneapolis, MN, 55455, USA; Department of Biochemistry, Molecular Biology & Biophysics, University of Minnesota, Minneapolis, Minnesota 55455, United States; Department of Molecular, Cellular, Developmental Biology, and Genetics

## Abstract

AAV is widely used for efficient delivery of DNA payloads. The extent to which the AAV capsid can be used to deliver a protein payload is unexplored. Here, we report engineered AAV capsids that directly package proteins – Protein Carrier AAV (pcAAV). Nanobodies inserted into the interior of the capsid mediate packaging of a cognate protein, including Green Fluorescent Protein (GFP), *Streptococcus pyogenes* Cas9, Cre recombinase, and the engineered peroxidase APEX2. We show that protein packaging efficiency is affected by the nanobody insertion position, the capsid protein isoform into which the nanobody is inserted, and the subcellular localization of the packaged protein during recombinant AAV capsid production; each of these factors can be rationally engineered to optimize protein packaging efficiency. We demonstrate that proteins packaged within pcAAV retain their enzymatic activity and that pcAAV can bind and enter the cell to deliver the protein payload. Establishing pcAAV as a protein delivery platform may expand the utility of AAV as a therapeutic and research tool.

## INTRODUCTION

Since its discovery as a contamination of Adenovirus productions in 1965 ^1,2^, studies of Adeno-associated virus (AAV) have spread the gamut from understanding its basic biology (reviewed in ^3,4^) to engineering AAV vectors for therapeutic applications (reviewed in ^5^). Being a non-enveloped virus belonging to the *Parvoviridae* family, AAV’s single-stranded DNA (ssDNA) genome spans 4.7 kb, is flanked by inverted terminal repeats (ITRs), and encodes two genes: *rep* and *cap* ^6^. While the *rep* gene encodes for several proteins mediating DNA replication, the *cap* gene encodes three viral proteins (VP1, VP2, and VP3) from the same open reading frame (ORF; ^7,8^), as well as the assembly-activating protein (AAP, ^9^) and membrane-associated accessory protein (MAAP, ^10^) from +1 ORFs. VP1, VP2, and VP3 form a 60-mer icosahedral capsid at an average stoichiometric ratio of 1:1:10, respectively (^11–13^). Assembly of the virions takes place in the nucleus, where the capsid is first formed and then the DNA loaded through the pore into the capsid ^14–16^.

For recombinant AAV production the natural genome can be replaced by an expression cassette of the transgene of interest, if the ssDNA cargo size does not exceed 4.7 kb ^17^. Multiple capsid engineering approaches, including peptide/domain insertions ^18–23^, DNA family shuffling ^24^, and recovery of ancestral AAV ^25^, have been established to modify the outside of the capsid with the goal to either enhance infection potency, refine cell tropism and/or enable immune evasion. Because of facile genome and capsid engineering, clinical success in several diseases ^26^, and a proven path to regulatory approval ^27^, AAV has emerged as a promising candidate for delivering therapeutic payloads.

The basic concept of AAV vectors for treating disease is to deliver a DNA sequence to target cells to modify the function of that cell directly (e.g., replace a faulty gene ^28–30^) or to reprogram that cell into an effector cell that corrects aberrant behaviors of other cells (e.g., generating CAR T cells ^31^). Aside from nucleic acids as a therapeutic agent, peptides and proteins (biologics) have garnered much attention as pharmaceuticals ^32^. Biologics come with unique delivery challenges, and several classes of nanoparticle delivery systems have been developed to overcome the limitations of free peptide/protein therapeutics ^33,34^. Encapsulation of peptide/protein biologics by nanoparticles has the potential to improve their stability, and enhance biodistribution and cellular uptake ^33^. The AAV capsid can be considered a type of self-assembling nanoparticle that has evolved to navigate and overcome environmental, systemic, and cellular delivery barriers. We thus wondered if the AAV capsid can be reengineered into a nanoparticle delivery vehicle for non-nucleic acid cargos, such as proteins.

In our previous broad domain insertion screen of the AAV-DJ capsid, we identified novel positions on the outside of the capsid into which domains, such as nanobodies, can be inserted to re-direct the tropism of the AAV. Surprisingly, the screening data also revealed sites in the capsid interior that can harbor domain insertions ^35^. While insertions at the inner surface are less useful with respect to optimizing cell tropism or immune evasion, we hypothesize that a domain insertion at the inside can be deployed to capture a cognate protein/ligand inside the capsid during AAV production, effectively converting AAV into Protein Carrier AAV (pcAAV).

To test this hypothesis, we inserted a GFP nanobody (GFPnb) into VP1 of AAV-DJ at the inside of the capsid. During AAV production we then replaced the AAV payload plasmid by a plasmid expressing GFP that lacked ITRs. Subsequent analysis of iodixanol gradient purified virus showed successful capturing of GFP protein instead of DNA within the AAV capsids. We were able to enhance the GFP protein packaging efficacy by inserting the nanobody into the more abundant VP3 and enriching the GFP in the nucleus during production. We demonstrate that AAV capsids packaging GFP protein can successfully deliver the payload to cells. Furthermore, we show that this novel protein packaging approach can be easily adapted to other proteins by substituting the GFPnb by an ALFA nanobody (ALFAnb), which in turn can bind any protein tagged with the short (13 amino acid) ALFA peptide. Using the two nanobodies, we demonstrate successful packaging of three proteins: *Streptococcus pyogenes* Cas9 (spCas9), Cre recombinase, and engineered ascorbate peroxidase (APEX2). Finally, we confirmed that a packaged enzyme – APEX2 in our case – retains its activity upon AAV packaging. To our knowledge, this is the first study showing that AAV can be transformed to a protein packaging vehicle, with implications for AAV biology and its use as a delivery vehicle for therapeutic payloads.

## RESULTS

### Engineering of the AAV capsid to package proteins

We hypothesized that the insertion of a binding domain into a position located on the interior of the capsid could enable the packaging of a cognate protein. For the binding domain, we chose to use a GFPnb, which is a single domain antibody that binds with high specificity and affinity (Kd = 1.4 nM) to GFP ^36,37^. We selected two positions in AAV-DJ VP1 located on the interior of the capsid to insert the GFPnb: H631 and H643. These positions were selected as they demonstrated minimal effects on AAV fitness in our previous domain insertion profiling of AAV-DJ VP1 ^35^. We mutated the start codons of VP2 (T138A) and VP3 (M203K, M211L, M235L) to ensure that the GFPnb is incorporated into only VP1. The VP1 containing the GFPnb was then supplemented with the wild type (wt) VP2/VP3, generated through mutation of the VP1 start codon (M1K), to enable capsid assembly (**Figure 1a**). We expect that the GFP protein will be packaged within AAV capsids through interaction with the GFPnb (**Figure 1b**), a process we predict to occur during capsid assembly (**Figure S1a**). To produce AAV packaging protein, we adapted the widely used helper-free AAV packaging protocol based on transient transfection of 293AAV cells ^38^. The two constructs containing the VP1 with the GFPnb insertion and wtVP2/VP3 were co-transfected with plasmids encoding a CAG-driven GFP (lacking ITRs), essentially replacing what would typically be a payload plasmid, and another encoding the necessary adenoviral helper genes. We subsequently purified the engineered AAV packaging protein using an iodixanol gradient removing any non-packaged GFP in the process. To determine if the binding domain was required for protein packaging, we also produced wtAAV-DJ in the presence of GFP expression.

**Figure 1:**
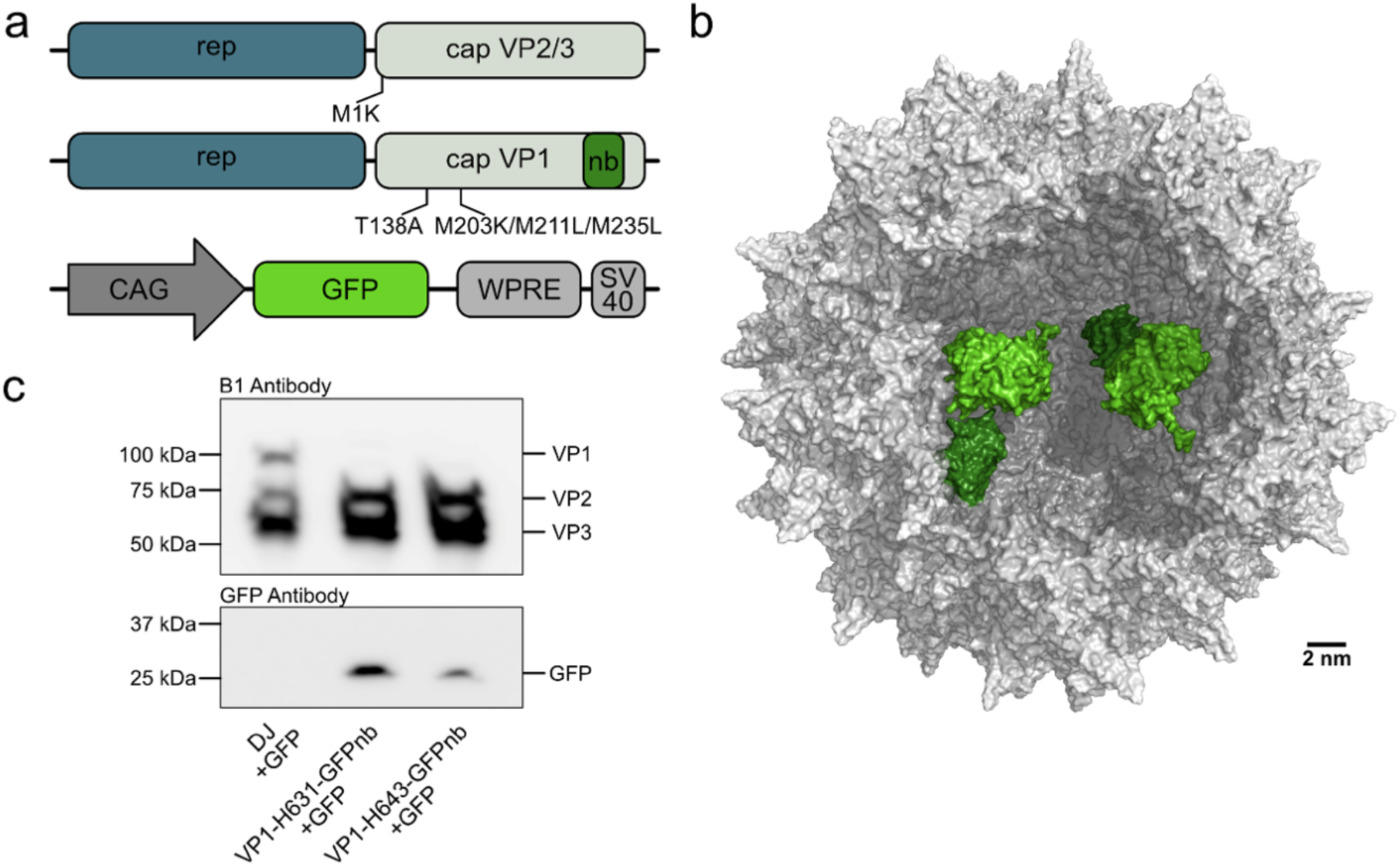
Insertion of a GFPnb into VP1 enables packaging of GFP within AAV capsids. **(a)** Schematic of the modified *cap* genes and GFP expression cassette used for protein packaging. **(b)** Model showing the AAV-DJ capsid (gray) containing two GFPnb (dark green) insertions at position H631 and packaging two GFP (light green). AAV-DJ capsid (RCSB PDB 7KFR); GFP:GFPnb complex (RCSB 3OGO). **(c)** Representative Western blot of wtAAV-DJ and AAV-DJ capsids incorporating VP1 with a GFPnb inserted at position H631 or H643 produced in cells overexpressing GFP during production.

Western blot analysis of the purified AAV using the B1 antibody, which recognizes an epitope common to VP1/VP2/VP3 ^16^, and a GFP antibody demonstrated successful packaging of GFP protein in capsids containing the GFPnb in VP1 at either H631 or H643. We detected no GFP protein in capsids lacking the binding domain, suggesting that the binding domain being present on the inside of the AAV capsid is required for packaging (**Figure 1c**). Both the degree of VP1 incorporation and GFP packaging efficiency differ based on the GFPnb insertion position with position H631 demonstrating higher VP1 and GFP signal than H643 (**Figure 1c; Figure S1b**). To verify that we do not have contamination of free GFP in the 40% iodixanol phase containing the assembled AAV capsids, we performed iodixanol gradient purification on cells expressing GFP protein and found that free GFP is located in the 25% phase (**Figure S2**). To confirm that the GFPnb was actually located in the interior of the capsid, we produced capsid variants with the GFPnb inserted at one of two positions: one located in the interior of the capsid (H631) and one located on the exterior of the capsid (T456). The localization of the GFPnb for each of these variants was determined using a pulldown assay specific to the GFPnb (**Figure S3a**). The capsid with the GFPnb inserted on the inside, position H631, demonstrated minimal enrichment in the pulldown assay compared to the capsid with the GFPnb inserted on the outside, position T456, consistent with the GFPnb being located within the capsid when inserted at position H631 (**Figure S3b**).

### Increasing efficacy of protein packaging

Our initial experiments demonstrated that the insertion of a GFPnb into VP1 at two different positions enabled the packaging of GFP protein into AAV-DJ capsids. We next sought to improve the efficiency of protein packaging through two complementary approaches: (1) increasing the concentration of the target protein in the nucleus, where capsid assembly occurs^16^, and (2) increasing the degree of incorporation of the binding domain containing subunit into assembled capsids. For the first approach, we added a c-Myc nuclear localization signal (NLS) to the C-terminus of the GFP protein (GFP-NLS). Western blot analysis of the iodixanol purified AAV demonstrated no conclusive increase in GFP packaging with the addition of the NLS (**Figure 2a**). While the addition of an NLS to the target protein demonstrated minimal benefits on protein packaging efficiency, we nevertheless decided to include the NLS in all subsequent protein packaging experiments. For the second approach, we inserted the GFPnb into VP3 instead of VP1, as VP3 is on average 10-fold more abundant in AAV capsids. Based on prior experience with nanobody insertions into VP3^23^, we reasoned that capsid assembly would be unlikely to occur without wtVP3 available and so we supplemented the VP3 with the GFPnb insertion with wtAAV-DJ VP1/VP2/VP3 on a separate construct in a 1:1 ratio (**Figure 2b**). wtAAV-DJ and AAV containing the GFPnb inserted into VP3 at either H631 or H643 were produced in cells expressing GFP-NLS followed by iodixanol purification. Subsequent Western blot analysis demonstrated an increase in protein packaging efficacy when compared to insertion of the GFPnb into VP1 (**Figure 2c; Figure S4**). The success of this approach suggests that our protein packaging system can be rationally engineered for improved packaging efficiency by considering known AAV biology, such as the location of capsid assembly and the principles behind the degree of subunit incorporation within the assembled capsid.

**Figure 2:**
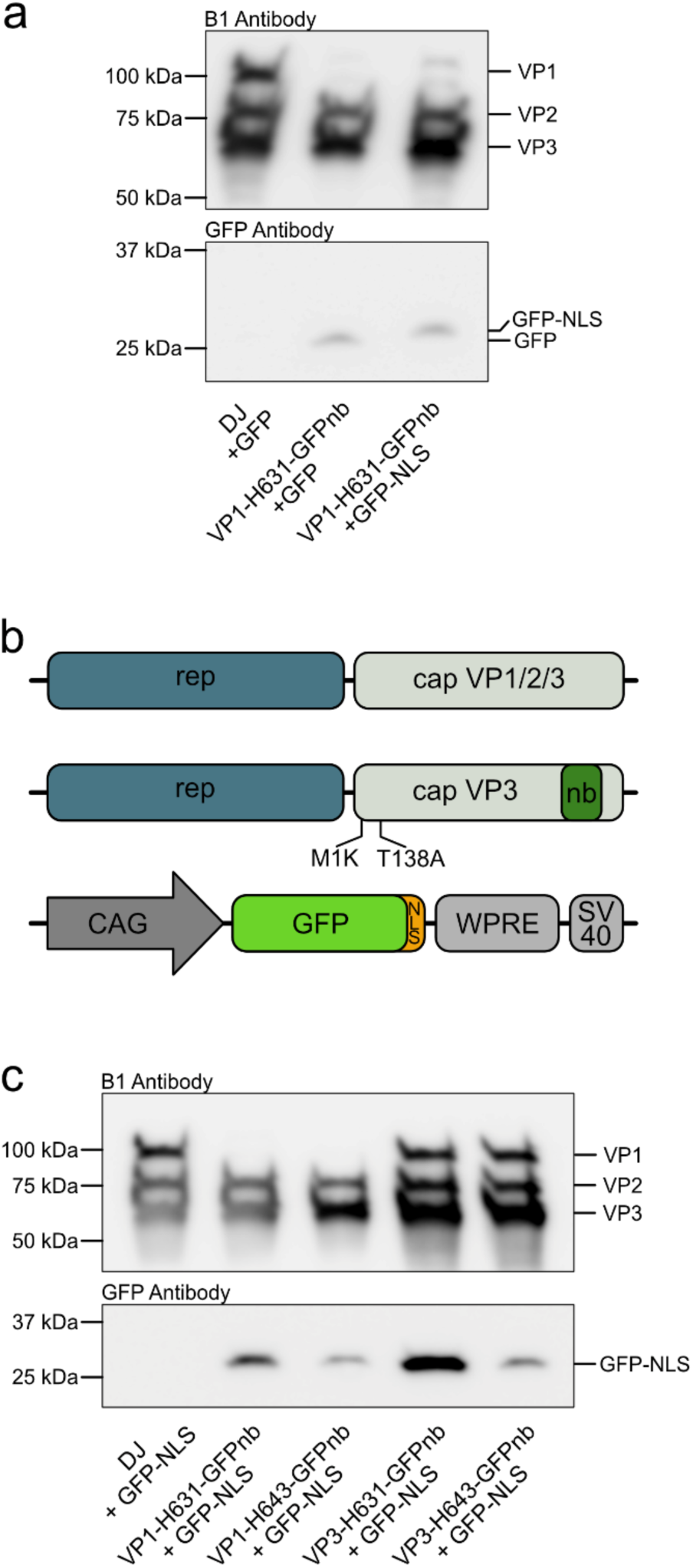
Addition of a c-Myc NLS to GFP and insertion of the GFPnb into VP3 increases protein packaging efficiency. **(a)** Representative Western blot comparing the packaging efficiency of wtAAV-DJ capsids and capsids incorporating a GFPnb in VP1 at position H631 at packaging GFP with or without the addition of a c-Myc NLS. **(b)** Schematic of the two *cap* genes used for producing AAV-DJ capsids with a GFPnb inserted into VP3 and the expression cassette used for generating GFP tagged with a c-Myc NLS at the C-terminus. **(c)** Representative Western blot comparing packaging of GFP protein tagged with a c-Myc NLS at the C-terminus in the indicated capsid variants. wtAAV-DJ produced in cells overexpressing GFP-NLS protein was included as a negative control.

### Quantification of protein packaging through label-free based mass spectrometry

Up to this point, we had been assessing the degree of protein packaging using a Western blot-based approach. To quantify how much target protein is packaged per capsid, we used label-free based mass spectrometry (MS). We produced and purified AAV containing a GFPnb inserted into either VP1 or VP3 at position H631. AAV capsids with the GFPnb inserted into VP1 were produced in cells overexpressing either GFP or GFP-NLS, while those with the GFPnb inserted into VP3 were produced in cells overexpressing GFP-NLS. These samples were analyzed by liquid chromatography with tandem mass spectrometry (LC MS/MS) using data-independent acquisition (DIA). The raw LC MS/MS data was used to calculate intensity based absolute quantification values (iBAQ). The iBAQ value is the sum of all peptide intensities divided by the number of theoretical peptides for a given protein and has been demonstrated to be a useful proxy of the relative molar abundance of a protein within a sample ^39,40^. The iBAQ values for GFP and VP proteins were used to calculate the ratio of GFP:capsid in each of the capsid variants (**Table 1**). For capsids with the GFPnb inserted into VP1 at position H631, we found ∼1 GFP per capsid and between 1-3 GFP-NLS per capsid. This aligns with our previous Western blot data showing a mild increase in GFP packaging with the addition of an NLS. Capsids with the GFPnb inserted into VP3 at position H631 packaged between 3-15 GFP-NLS per capsid consistent with a significant increase in protein packaging efficacy when the GFPnb is inserted into VP3 instead of VP1.

**Table 1:** Mass spec quantification of select AAV capsids packaging GFP protein variants. Data from three biological replicates are shown.

| VP insertion | Nanobody | Payload | GFP:capsid |  |  |  |
| --- | --- | --- | --- | --- | --- | --- |
|  |  |  | Rep1 | Rep2 | Rep3 | Avg |
| WT | - | GFP-NLS | 0.02 | 0.17 | 0.06 | 0.08±0.06 |
| VP1 | GFPnb | GFP | 0.89 | 0.73 | 0.33 | 0.65±0.24 |
| VP1 | GFPnb | GFP-NLS | 0.89 | 2.27 | 0.81 | 1.33±0.67 |
| VP3 | GFPnb | GFP-NLS | 13.14 | 2.17 | 1.35 | 5.55±5.37 |
| VP3 | ALFAnb | GFP-NLS-ALFA | 8.63 | 1.11 | 0.30 | 3.35±3.75 |

### Delivering an AAV protein payload to cells

We next determined if our engineered capsids retained the ability to infect and deliver a protein payload to cells. 293AAV cells were transduced with capsids containing the GFPnb inserted into VP3 at position H631 and packaging GFP-NLS as this capsid variant previously demonstrated the highest protein packaging efficacy (**Figure 2c**, **Table 1**). As negative controls, we included cells alone, cells transduced with wtAAV-DJ produced in cells overexpressing GFP-NLS, and cells incubated with 1E5 molecules recombinant GFP per cell. Live cell fluorescence microscopy was performed ten hours post transduction to assess the delivery of GFP protein to cells. The nucleus and cell membrane were counterstained to facilitate the localization of the delivered protein. As seen in **Figure 3a**, GFP signal was detected exclusively in cells treated with AAV packaging GFP-NLS protein. The GFP signal was distributed in puncta throughout the cytoplasm of the cells. We saw little to no GFP signal located within the nucleus, which aligns with previous reports that AAV capsids lacking a genome are not transported to the nucleus ^41^. An automated image processing pipeline was subsequently used to quantify the mean GFP fluorescence signal for each sample (**Figure 3b, Figure S5**). We found that cells treated with AAV packaging GFP-NLS protein demonstrated a significant increase in mean GFP intensity when compared to cells alone. These results establish that engineered Protein Carrier AAV capsids can deliver a protein payload to cells.

**Figure 3:**
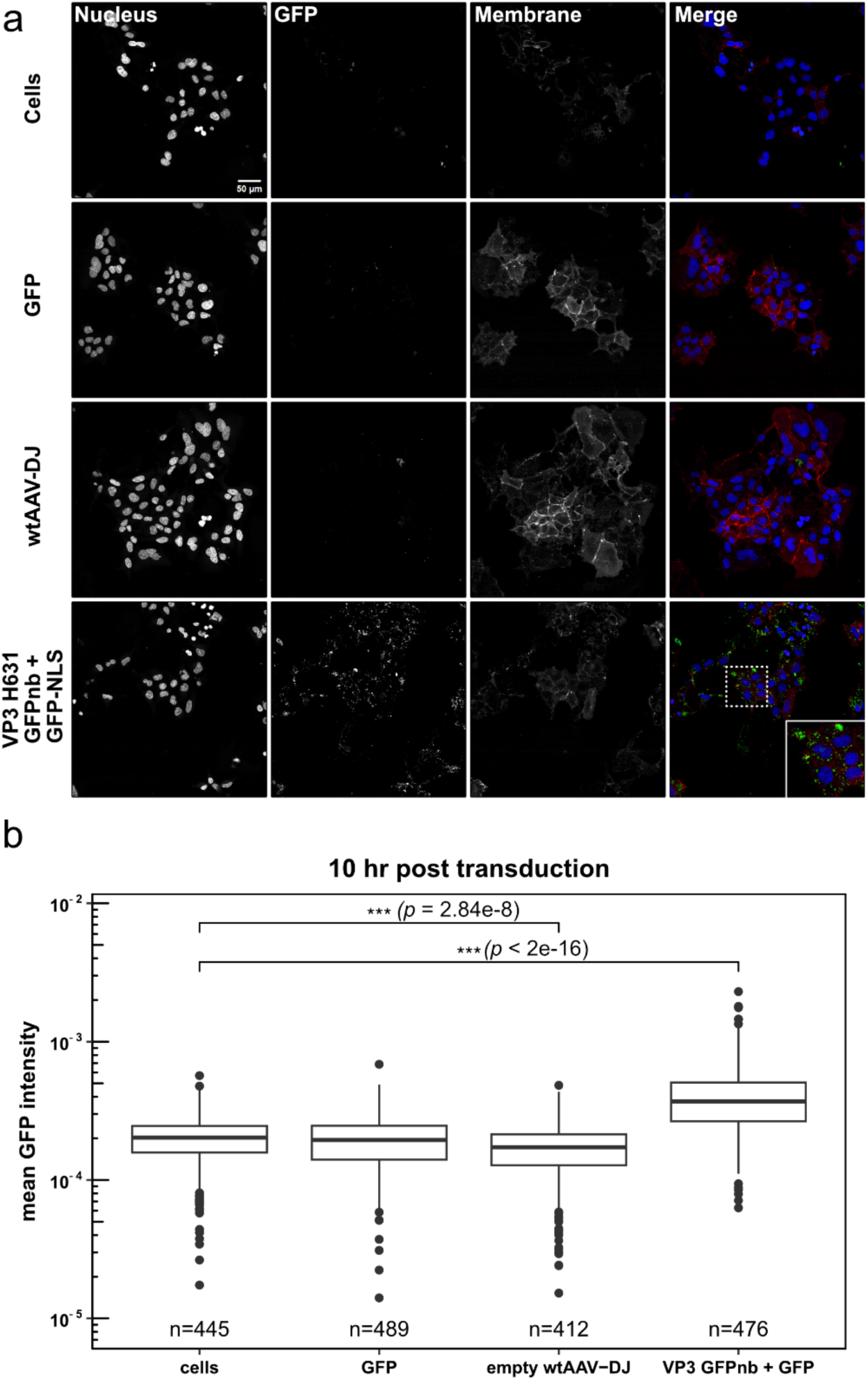
Live cell imaging of AAV mediated GFP protein delivery. **(a)** The delivery of GFP protein by AAV capsids incorporating a GFPnb in VP3 at position H631 was visualized using fluorescence microscopy. 24 hours post seeding, 293AAV cells were either mock transduced, treated with 1E5 molecules recombinant GFP-ALFA protein per cell, or incubated with 15 μL of either wtAAV-DJ or AAV-DJ capsids incorporating a GFPnb at position H631 in VP3 and packaging GFP-NLS protein. Ten hours post transduction, the nucleus and cell membrane were labeled and subsequently imaged. Representative images from at least four fields of view. **(b)** An automated image processing pipeline was used to quantify the GFP fluorescence intensity for each of our treatment conditions. The log transformed mean GFP fluorescence intensity values are represented in a box plot, with the number of cells quantified in each condition indicated in the figure. The box represents the interquartile range (IQR), where the horizontal bar represents the median and the top and bottom hinges represent the 25th and 75th percentiles, respectively. Whiskers extend to ±1.5 times the IQR with points representing potential outliers. \*\*\**p* < 0.001 by Kruskal-Willis test followed by a post hoc Dunn’s test with Benjamini-Hochberg correction.

### Packaging spCas9 protein

Next, we determined the ability of our engineered AAV capsid to package proteins other than GFP. The first protein we packaged was the gene editing tool spCas9 tagged with a GFP at the C-terminus (spCas9-GFP). AAV capsids incorporating a GFPnb inserted into VP1 at H643 were produced in cells co-transfected with a spCas9-GFP construct or with a spCas9 construct lacking the GFP fusion protein. wtAAV-DJ was produced in cells simultaneously transfected with the spCas9-GFP construct as a negative control. Surprisingly, Western blot analysis of the iodixanol purified virus utilizing B1 and Cas9 antibodies demonstrated that packaging of spCas9-GFP occurred both in capsids with the GFPnb insertion and in the wtAAV-DJ negative control. Additionally, packaging of spCas9 lacking the GFP tag was seen in capsids with the GFPnb insertion (**Figure 4a**). These data suggest that nonspecific packaging – which we define as packaging of the target protein in the absence of either the binding domain or its cognate ligand – occurs frequently for spCas9. Considering that nonspecific packaging did not occur when packaging GFP, differences in the subcellular localization and biochemical properties of the target protein may affect its propensity to be packaged either specifically or nonspecifically. To test the ability of spCas9 protein packaged within AAV to induce editing, we co-transduced cells with AAV containing a GFPnb inserted into VP1 at position H643 and packaging spCas9-GFP protein with an AAV containing a DNA genome encoding for a guide RNA targeting a test locus. However, we were unable to detect any degree of editing at the target locus (data not shown). We speculate that this is related to the defective nuclear trafficking, and therefore nuclear delivery, of Protein Carrier AAV.

**Figure 4:**
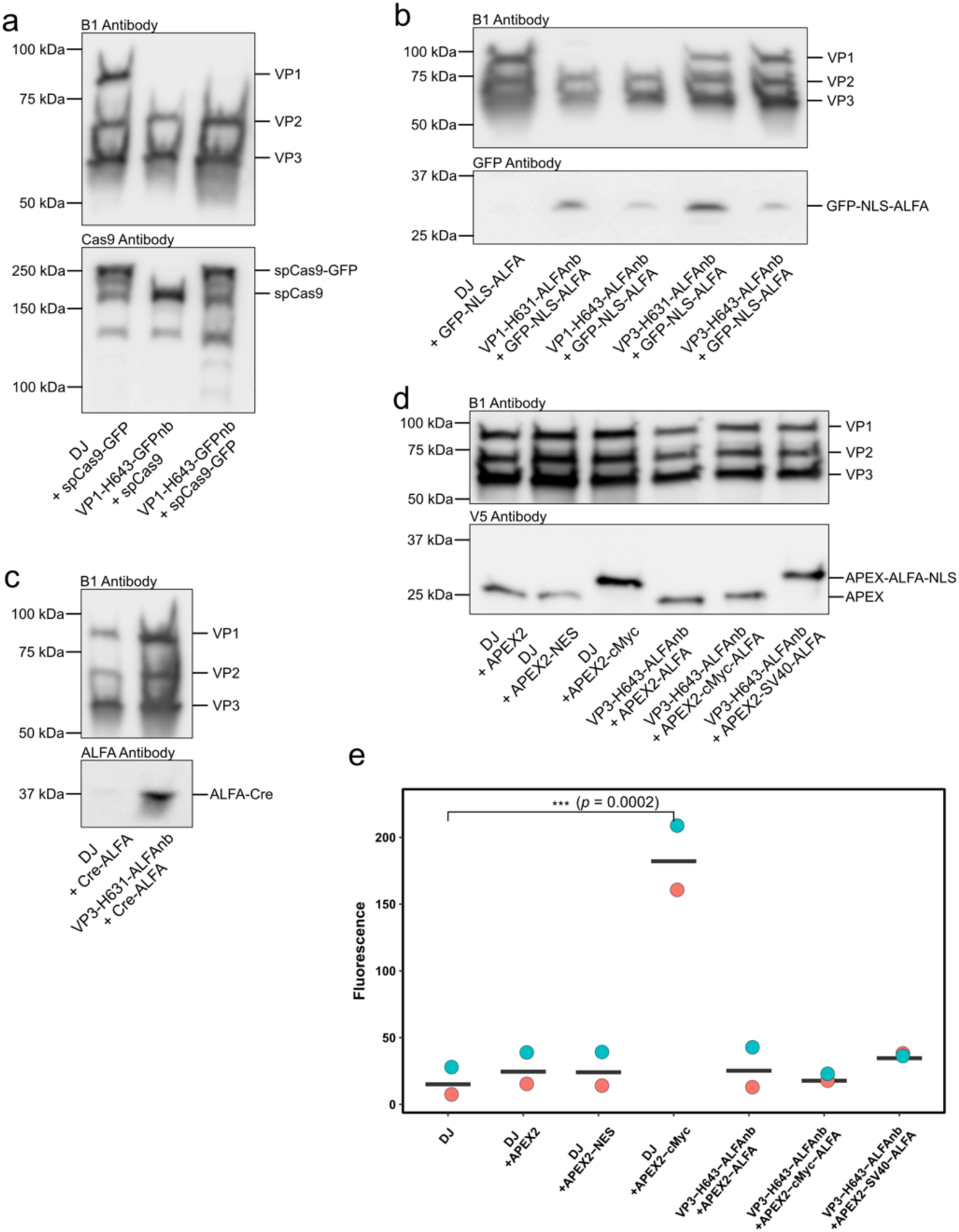
Engineered AAV capsids can package several proteins, which can retain enzymatic activity. **(a)** Representative Western blot demonstrating packaging of spCas9 protein with or without a GFP tag in wtAAV-DJ capsids or capsids with a GFPnb inserted into VP1 at position H643. The B1 antibody and a spCas9 antibody were used for visualizing the VP subunits and spCas9, respectively. **(b)** Representative Western blot demonstrating that GFP protein tagged with both a c-Myc NLS and an ALFA tag is packaged within capsids containing the ALFAnb inserted into either VP1 or VP3 at either position H631 or H643. The B1 antibody and a GFP antibody were used for visualizing the VP subunits and GFP, respectively. **(c)** Representative Western blot showing packaging of Cre recombinase protein tagged with an ALFA tag. The B1 antibody and an ALFA tag antibody were used for visualizing the VP subunits and ALFA tagged Cre, respectively. **(d)** Representative Western blot demonstrating packaging of the indicated APEX2 constructs in both engineered and wtAAV-DJ capsids. The B1 antibody and V5 antibody were used for visualizing the VP subunits and V5 tagged APEX2, respectively. **(e)** Fluorescence values per 1 µL of virus obtained for each AAV-DJ capsid variant packaging the indicated APEX2 constructs. Points represent biological replicates with lines indicating the mean for each sample. Biological replicate one is colored blue and replicate two is colored orange. \*\*\**p* < 0.001 by Dunnett’s test.

### Packaging of Cre and APEX2 using the ALFAnb

The use of the GFPnb as the binding domain requires tagging a target protein with the 27 kDa GFP protein to enable packaging within our engineered capsids. To expand the versatility of our protein packaging system, we explored changing the binding domain to the ALFAnb (**Figure S6**). The ALFAnb binds with high specificity and affinity (∼26 pM) to a rationally designed 13 amino acid sequence, the ALFA tag, that forms a neutrally charged, hydrophilic, alpha helix ^42^. The ALFA tag, being both small and having little to no effect on the function of the tagged protein, is an ideal fusion tag for our protein packaging system. We first inserted the ALFAnb into either VP1 or VP3 at H631 or H643 and tested the ability of capsids incorporating the nanobody containing subunit to package GFP C-terminally tagged with a c-Myc NLS followed by the ALFA tag (GFP-NLS-ALFA). Western blot analysis demonstrated successful packaging of GFP-NLS-ALFA protein in all capsids incorporating an ALFAnb containing subunit, while no packaging was seen in the wtAAV-DJ negative control (**Figure 4b; Figure S7**). Like our results using the GFPnb, the degree of protein packaging was dependent on both the insertion position and the identity of the VP containing the insertion. Accordingly, insertion into VP3 at position H631 demonstrating the highest protein packaging efficacy.

To better compare the packaging efficiency of capsids incorporating the ALFAnb versus the GFPnb, we used our LC MS/MS approach to determine the GFP:capsid ratio of capsids incorporating VP3 with the GFPnb at position H631 and packaging GFP-NLS-ALFA protein. These capsids were found to package between 1-9 GFP-NLS-ALFA protein per capsid, while the comparable GFPnb incorporating capsids packaged between 3-15 GFP-NLS-ALFA protein per capsid (**Table 1**). This suggests that the ALFAnb is slightly less efficient than the GFPnb at mediating protein packaging.

Having established the ALFAnb as an alternative binding domain for protein packaging, we selected two target proteins for ALFA tag mediated protein packaging into AAV capsids: the recombinase Cre and the peroxidase APEX2. Cre recombinase is a widely used tool for genetic manipulation as it catalyzes the site-specific recombination of DNA through the recognition of specific nucleotide sequences termed *loxP* sites ^43^. We produced wtAAV-DJ and capsids incorporating VP3 with the ALFAnb inserted at position H631 in cells co-transfected with a Cre recombinase fused to an ALFA tag and SV40 NLS at the C-terminus (Cre-ALFA). Western blot analysis demonstrated successful packaging of Cre-ALFA protein in capsids incorporating the ALFAnb (**Figure 4c**). A faint band is visible in the wtAAV-DJ negative control suggesting that Cre-ALFA protein, similarly to spCas9, undergoes some degree of nonspecific packaging. We subsequently performed a Cre recombinase activity assay to determine if AAV mediated delivery of Cre protein could catalyze recombination, however, no recombinase activity was observed (data not shown), again likely due to lack of nuclear trafficking.

As a second target protein, we chose to package the heme peroxidase APEX2, which has applications in labelling intracellular proteins for electron microscopy (EM) and proximity tagging of proteins with biotin for spatiotemporal proteomics ^44,45^. We produced wtAAV-DJ and capsids incorporating VP3 with the ALFAnb inserted at position H631 in cells simultaneously transfected with either APEX2 alone or C-terminally tagged with one of the following: (1) a c-Myc NLS, (2) a nuclear export signal (NES), (3) a c-Myc NLS and ALFA tag, or (4) a SV40 NLS and ALFA tag. Interestingly, the Western blot demonstrated packaging of all variants of APEX2 irrespective of ALFAnb incorporation in the capsid (**Figure 4d**). In fact, incorporation of the ALFAnb, to capture ALFA-tagged APEX2, did not increase APEX2 content within AAV capsids. Addition of a NES decreased APEX2 packaging somewhat, and nuclear localization of APEX2 (by addition of an NLS) appeared sufficient for efficient packaging. The nonspecific packaging seen here was similar to that observed for spCas9 further underscoring that the biochemical properties of the target protein may affect packaging efficiency.

### Packaged APEX2 protein retains enzymatic activity

In light of inconclusive results for SpCas9 and Cre, we asked if proteins retain enzymatic activity when packaged into Protein Carrier AAV. An *in vitro* peroxidase assay that produces a fluorescent readout was used to test the activity of each of the purified AAV variants packaging APEX2 described above. The resulting fluorescent values were normalized to wtAAV-DJ not produced in the presence of APEX2. We found that wtAAV-DJ packaging APEX2-cMyc showed significantly higher fluorescence compared to the empty wtAAV-DJ control (**Figure 4e**). These experiments establish that an enzyme packaged within AAV capsids can retain its activity.

## DISCUSSION

AAV has achieved success in both research and therapy as it can efficiently deliver a DNA payload *in vitro* and *in vivo*. The efficiency by which AAV delivers DNA to a target cell is due to evolved functions of the AAV capsid, which mediate each step of the infectious pathway including cell surface binding, endosomal escape, and nuclear transport ^46–49^. Each of these characteristics are required for the delivery of protein cargos to cells, but are difficult to engineer into new bottom-up designed synthetic delivery systems, such as lipid nanoparticles ^50,51^. Alternative approaches that overcome biological barriers to delivery of therapeutic cargos, including protein therapeutics, are needed. Here, we take the initial steps towards leveraging naturally evolved characteristics of the AAV capsid for the packaging and delivery of protein payloads.

To our knowledge, this study represents the first instance of packaging proteins into AAV capsids and establishes some of the basic principles that determine the efficiency of protein packaging. On the capsid side, the position of the binding domain insertion and the identity of the VP into which it was inserted both affected the efficiency of protein packaging. The two binding domain insertion positions, H631 and H643, were selected based on their permissibility to be inserted into the capsid, while minimally affecting the DNA packaging and infectivity of the resulting virus ^35^. Both insertion positions are located beneath the 3-fold protrusion with residue H631 being a part of a nucleotide binding pocket of the capsid that is highly conserved across multiple AAV serotypes ^52–55^. Residue H643 lies just outside of this nucleotide binding pocket and is also conserved amongst multiple AAV serotypes (**Figure S8**). Insertion of the binding domain at position H631 consistently produced higher packaging efficacy when compared to insertion at H643. This suggests that exploring additional insertion positions for the binding domain has the potential to further increase packaging efficiency. Similarly, the increase in protein packaging seen when inserting the binding domain into VP3 instead of VP1 suggests optimization of the incorporation ratio of the domain containing subunit could further improve protein packaging. The fact that both binding domain insertion positions are conserved suggests that these positions could be used to adapt this approach for generating pcAAV to additional AAV serotypes, thus unlocking AAV-mediated tissue-specific delivery of protein payloads. The extension of our protein packaging system to additional AAV serotypes would also provide an opportunity to study how differences in capsid assembly including changes in the localization of assembling capsids (e.g., nucleus versus nucleolus) ^56^ and AAP dependence or independence ^57^ affects protein packaging.

Both protein packaging efficiency and nonspecific protein packaging are affected by characteristics of the target protein with subcellular localization playing a large role. We hypothesize that the process of protein packaging within AAV occurs during the capsid assembly process, during which the target protein associates with the inserted binding domain and becomes packaged within the assembled capsid. We predicted that the inclusion of an NLS on the target protein could improve packaging efficiency through increasing the localization of the target protein at the site of capsid assembly. However, we saw only modest differences in protein packaging for GFP or APEX2 with the addition of an NLS (**Figure 2a**, **Figure 4d**). Previous studies have shown that proteins with a molecular mass below 60 kDa passively diffuse into the nucleus. Larger proteins have also been demonstrated to accumulate in the nucleus through passive diffusion albeit at a significantly reduced rate ^58^. In the instance of the smaller proteins packaged (GFP, APEX2, and Cre), it is likely that the passive diffusion of protein to the nucleus provides sufficient protein abundance for packaging. The high degree of nonspecific packaging of spCas9 protein into capsids lacking a binding domain further supports the idea that protein packaging is highly dependent on subcellular localization. In particular, spCas9 contains a nucleolus detention signal (NoDS) that drives strong localization to the nucleolus ^59^. As discussed previously, the nucleolus is the site of capsid assembly for most AAV serotypes ^16,56^, thus suggesting that the inclusion of a NoDS in future target proteins could be a potential strategy to increase protein packaging efficiency. Previous studies have demonstrated that nucleolar-associated proteins (e.g., nucleolin and nucleophosmin) are common contaminants in AAV productions and interact with the capsid during both production and infection ^60–62^. Our data suggests that packaging of nucleolar enriched proteins within the capsid may deserve further attention as a possible source for contaminant host proteins in AAV productions.

Our live cell imaging experiments demonstrated that AAV capsids packaging protein do not deliver the protein payload to the nucleus, which currently represents a significant challenge to the utility of Protein Carrier AAV. Johnson et.al. showed that empty AAV2 capsids are not transported to the nucleus, however, the exact reasons behind this are poorly understood ^41^. It is possible that AAV requires the presence of a DNA genome to trigger the externalization of VP1u, since revealing the phospholipase domain is essential for endosomal escape and the exposed NLS for subsequent nuclear transport. Alternatively, it is possible that protein packaging prevents VP1u internalization. Previous studies have indicated that preexposure of VP1u leads to significantly reduced infectivity, which suggests that infection potency is sensitive to the temporal and spatial regulation of VP1u exposure ^63^. Our trafficking data provides a potential explanation as to why both our spCas9 and Cre activity assays exhibited no activity. It is likely that both proteins remained trapped within the AAV capsid during intracellular trafficking and so were destined to the same fate as empty capsids – degradation in the lysosome. Our activity assay with APEX2 demonstrates that enzymes packaged within AAV capsids retain functionality meaning that if we can rescue intracellular trafficking of AAV capsids packaging proteins, it is likely that the delivered protein will remain functional. Future research clearly should be directed to rescuing nuclear trafficking of Protein Carrier AAV as this technology has the potential to be broadly useful for transient delivery of protein cargo. In the field of gene editing, delivering gene editing tools as proteins carries several benefits. For example, the delivery of spCas9 as a protein or as a ribonucleoprotein (RNP), wherein the spCas9 protein is in complex with the guide RNA, increases editing efficiencies, while decreasing off target editing and genotoxicity ^64–66^.

Extensive work has been performed to generate novel capsid variants with highly specific tissue tropism ^20,67–69^. Extension of protein packaging to these capsid variants could enable the tissue-specific delivery of proteins in difficult-to-reach tissues, such as the brain, without the need for validating extensive retargeting strategies. Conservation of binding domain insertion positions across a wide range of AAV serotypes, suggest that adapting our protein packaging strategy to these serotypes would be achievable. Additionally, our engineered AAV capsids could be used to deliver potentially any protein of interest that falls within the size limitations of the capsid, which we demonstrated can at least accommodate proteins up to a size of 194.2 kDa, the size of the spCas9. Taken together, expanding the natural payload of AAV beyond DNA to protein may significantly increase the utility of an already versatile viral vector.

## MATERIAL AND METHODS

### Cloning

The GFP expression plasmid was generated by amplifying the entire expression cassette from pAAV-CAG-GFP (Addgene #37825) and subcloning it into pATT-Dest (Addgene plasmid #79770). For the GFP-NLS plasmid a c-myc NLS was added to the C-terminus of the GFP by Gibson Assembly. Similarly, the GFP-NLS-ALFAtag plasmid was assembled by inserting a gBlock encoding the c-Myc NLS and the ALFAtag separated by GS-linker. The plasmid pdCas9-GFP was obtained from Addgene (Addgene #181906) and the point mutations D10A and H840A were reversed to obtain spCas9-GFP. The plasmid spCas9 was generated by removing the coding sequence of GFP of the aforementioned spCas9-GFP plasmid and adding a stop codon behind the spCas9 coding sequence. The Cre-ALFA plasmid was generated by replacing the Cas9-GFP coding sequence by a gBlock encoding an ALFA-SV40-NLS-Cre coding sequence using the restriction sites NheI and NotI. The ALFA tag is separated by a GGGGS-linker from the SV40-NLS. The APEX2 expression plasmid was obtained from APEX2-NLS (Addgene #124617). The NLS was removed or replaced with a c-Myc NLS to generate the APEX2 only and APEX2-cMyc-NLS plasmids respectively. Separately, the SV40-NLS was codon optimized, ordered as a gBlock, and used to generate a APEX2-SV40-NLS plasmid. Two oligos encoding the ALFA tag were annealed and used to insert the ALFA tag behind the SV40-NLS to generate the final APEX2-SV40-NLS-ALFA plasmid. A gBlock containing the ALFA tag was inserted to generate the APEX2-cMyc-NLS-ALFA plasmids, while a gBlock containing the nuclear export signal (NES) was inserted to generate the APEX2-NES plasmid. All cloning for the APEX2 constructs was done using Gibson assembly. The DJ-M1K plasmid as well as DJ-VP1 and DJ-VP3 were previously published by us ^35,70^. The GFPnb or ALFAnb were inserted behind the residues H631 or H643 by Golden Gate assembly ^71^ using BsmBI overhangs. The nanobodies were flanked by SGGGG linkers.

All plasmids used or generated in this study are listed in **Table S1**. Select GFP, spCas9, Cre, and APEX2 protein sequences are shown in **Figure S9**. Restriction enzymes for cloning were obtained from New England Biolabs (NEB), oligos and gBlocks from Integrated DNA Technologies (IDT), and plasmid were isolated using either the Zyppy Plasmid Miniprep Kit, ZymoPURE II Midiprep Kit or the ZymoPURE II Maxiprep Kit all from Zymo Research. For all PCR amplifications the PrimeSTAR Max DNA Polymerase (Takara Bio Inc.) was used by following the manufacturer’s instructions. PCR products were analyzed on 1 % agarose gel and purified using the Zymoclean Gel DNA Extraction Kit (Zymo Research). Transformations were made into NEB Stable Competent E. coli (Thermo Scientific) and cells plated on LB plates containing 100 µg/mL carbenicillin.

### Tissue culture

293AAV cells (Cell Biolabs) and 293FT cells (Invitrogen) were cultured in DMEM (Gibco) containing 4.5 g/L D-glucose, L-glutamine, 110 mg/L sodium pyruvate, and supplemented with 10 % fetal bovine serum (Gibco) and 100 U per mL penicillin/100 μg per mL streptomycin (Gibco). Cells were kept in a humidified cell culture incubator at 5 % CO2 and 37 °C and passaged every 2-3 days when reaching 70-90 % confluency.

### Iodixanol gradient AAV purification

AAV were purified using iodixanol gradients according to previously published protocols ^72,73^. In brief, five million 293AAV cells were seeded into 15 cm dishes. 48 hours post seeding, cells were transfected with 47 µg DNA per dish using PEI. The DNA was split up in an equimolar ratio between (i) an Adeno-helper plasmid, (ii) a cargo plasmid encoding either a transgene flanked by ITR for ssDNA packaging or a transgene without ITRs for protein packaging, (iii) a plasmid encoding the *rep* gene and *cap* gene encoding for a VP with the nanobody insertion, and (iv) a plasmid complementing the remaining VP proteins. Cells from ten dishes each were harvested 72 hours post transfection, washed with PBS once and resuspended in a buffer containing 2 mM MgCl2, 0.15 M NaCl and 50 mM Tris-HCl at pH 8.5. Next, five freeze and thaw cycles were performed by alternating between liquid nitrogen and a 37 °C water bath to lyse the cells and free the AAV particles. Free plasmid and genomic DNA was digested with Benzonase Nuclease (Sigma-Aldrich) for one hour at 37 °C. To separate the cell debris from the AAV lysate, samples were spun down twice at 4,000 xg at 4 °C. The AAV lysate was then loaded into ultracentrifugation tubes (Beckman Coulter) and iodixanol (Iodixanol-OptiPrepTM, Progen) of 15 %, 25 %, 40 %, and 60 % was layered underneath. The density gradient centrifugation was performed in a 70.1Ti rotor (Beckman Coulter) for 2 hours at 50,000 rpm and at 4 °C. Post centrifugation, the 40 % phase containing the AAV particles was isolated. For the Western blot analysis of Figure S2 fractions of the 60 %, 40 %, and 25 % phase were taken instead of just the entire 40 % iodixanol phase.

### qPCR

To assess titers of gradient purified AAV carrying a DNA payload qPCR was performed as follows: 1 µl of purified AAV were mixed with 43µl PBS, 5 µl Proteinase K buffer (100 nM Tris-HCl, pH 8.0, 10 mM EDTA, and 10 % SDS) and 1 µl Proteinase K (20 mg/ml; Zymo Research). Subsequently, samples were incubated for 30 min at 50 °C to free the ssDNA payload, followed by heat inactivation of the Proteinase K for 10 min at 95°C. The ssDNA was purified using the DNA Clean & Concentrator-5 Kit (Zymo Research) kit according to the manufacturer’s instructions and samples diluted 1:1,000. qPCR was done with the PowerUp SYBR Green Master Mix (Applied Biosystem) on a QuantStudio5 Real-Time PCR System (Applied Biosystems). A primer set binding in the CMV-enhancer ^74^ (forward: AACGCCAATAGGGACTTTCC, reverse: GGGCGTACTTGGCATATGAT) of the ssDNA payload was used together with a plasmid standard at a known concentration to calculate the titer in vg/ml.

### Western Blot

30 µl of purified AAV were mixed with 10 µl 4x Laemmli Sample Buffer (Bio-Rad), supplemented with 10% 2-mercaptoethanol and denatured for 10 min at 95 °C. Samples were then separated by molecular weight on a 4-20 % precast polyacrylamide gel (Bio-Rad), alongside 7.5 µl of the Precision Plus Protein Dual Color Standard (Bio-Rad), for 90 min at 120 V in in Tris/Glycine/SDS Electrophoresis Buffer (Bio-Rad). Subsequently, proteins were transferred onto a nitrocellulose membrane (pore size 0.45 µm; Thermo Scientific) in an ice-cold blotting buffer (25 mM Tris Base, 96 mM glycine, 20 % methanol) for 80 minutes at 110 V. Post blotting, the membrane was washed once for 2 min in TBS-T buffer (20 mM Tris Base, 137 mM NaCl, pH 7.6, 0.05% Tween-20) and then incubated in blocking solution (5 % skim milk in TBS-T) for 1 h at room temperature on a rocker. Before the primary antibody was added, the membrane was then cut horizontally at ∼45 kDa. The upper half received a primary antibody binding all three VP proteins (1:250; anti-AAV VP1/VP2/VP3 mouse monoclonal, B1, supernatant, Progen), whereas the lower half was incubated in a primary antibody binding GFP (1:1,000; anti-GFP mouse monoclonal, Invitrogen MA5-15349). Both primary antibodies were diluted in 5 % skim milk in TBS-T and incubated overnight at 4 °C. The next day, the membrane was washed four times for 5 min in TBS-T before the secondary antibody was added (1:50,000 in 5% skim milk in TBS-T; anti-mouse IgG-peroxidase antibody produced in goat, Sigma-Aldrich). The secondary antibody was incubated for an hour at room temperature on a rocker. Afterwards, the membranes were washed again four times for 5 min in TBS-T and subsequently, the SuperSignal West Dura Extended Duration Substrate kit solution (Thermo Scientific) was applied. The chemiluminescence signal was detected with an Amersham Imager 600 (GE Healthcare). Quantification of VP expression was done using ImageJ. All uncropped Western blots are available in **Figure S10**.

### Live cell imaging

293AAV cells were seeded into a 96 well glass bottom plate (Greiner Bio-One) at a density of 12,000 cells/well in 100 μL culture media. 24 hours following cell seeding, the media was aspirated, and cells were incubated with either 15 μL of iodixanol purified virus or 1E5 molecules per cell of recombinant GFP-ALFA protein (NanoTag Biotechnologies) diluted up to 100 μL in culture media. The media of mock transfected cells was replaced at this time. 10 hours following the treatment cells were prepared for live cell imaging through first staining with a WGA Alexa Fluor 555 conjugate (Invitrogen W32464) then with NucBlue Live ReadyProbes Reagant (Invitrogen R37605) in the following manner. The culture media was removed from all samples then replaced with 50 μL of a 1.0 mg/mL WGA Alexa Fluor 555 conjugate solution prepared in Hanks Balanced Salt Solution (HBSS). Samples were incubated in the labeling solution at 37 °C for 10 minutes then washed twice with 100 μL of prewarmed (37°C) FluoroBrite DMEM (Gibco). Following the last wash, 100 μL of warm FluoroBrite DMEM was added to each sample. Immediately prior to imaging, 12 μL of NucBlue Live ReadyProbes Reagant was added to each sample and incubated at room temperature for 20 minutes.

All samples were imaged using a Nikon A1Rsi HD Confocal with SIM Super Resolution using a Nikon PlanApo VC 20x 0.75 NA objective with acquisition being performed using Nikon Elements. Nuclear Hoechst 33342 (NucBlue Live ReadyProbes) labeling was excited with a 405 nm laser and emission selected with a 460/50nm filter. AAV delivered enhanced GFP (EGFP) was excited with a 488 nm laser and emission selected with a 525/50nm filter. WGA Alexa Fluor 555 cell membrane labeling was excited with a 561 nm laser and emission selected with a 620/60nm filter. The 96-well glass bottom plate was placed onto a Tokai Hit Stage Top Incubator (model no. INUB-GSI-BP-F1) and maintained at 37 °C and 5 % CO2 for the duration of imaging. Galvano scanning mode was used to acquire at least four unique fields of view for each sample. Images were then processed for automated imaging analysis in the following manner. First, spectral overlap between the WGA Alexa Fluor 555 labeling and EGFP required the subtraction of the WGA 555 channel from the EGFP channel which was achieved using a custom-built FIJI (version 1.54f) ^75^ macro. The processed images were then used as input for a CellProfiler (version 4.2.6) ^76^ pipeline (**Figure S5**). Briefly, nuclei were identified as primary objects through the DAPI channel using a global thresholding strategy and an Otsu thresholding method. Next, cells were identified as secondary objects, using nuclei as input objects and the WGA 555 channel as the input image, through a global thresholding strategy and a Minimum Cross-Entropy thresholding method. From here, the GFP fluorescence intensity was measured for each object identified as a cell and subsequently exported as a spreadsheet. The output measurements were subsequently analyzed using a custom-built R-script (R version 4.3.3) to determine the mean GFP fluorescence intensity for each sample.

### Mass spectrometry

All AAV samples submitted for mass spectrometry analysis were produced from ten 15cm dishes of 293AAV cells then purified by iodixanol gradient ultracentrifugation as described above. The iodixanol purified AAV were subsequently buffer exchanged to PBS using a 10 kDa MWCO filter (Amicon) in the following manner. All buffers were used ice cold and all centrifugation steps were performed at 3,140 xg and 4 °C. The 10 kDa MWCO filter was first equilibrated by twice adding PBS to the max fill line and centrifuging until the level of PBS was at the level of the filter. 1 mL of iodixanol purified AAV was then added to the filter and then topped up to the max fill line with PBS. The sample was mixed via pipetting until the solution appeared homogenous. The sample was centrifuged until the solution was at the level of the filter at which point PBS was added to the max fill line and the sample was mixed via pipetting until it was homogenous. These steps were repeated until no iodixanol could be seen remaining in the solution at which point the sample was concentrated via centrifugation down to 250 μL. The concentrated samples were submitted to the University of Minnesota Center for Metabolomics and Proteomics where LC MS/MS analyses were performed on an Orbitrap Fusion Tribrid (ThermoFisher Scientific) equipped with a ThermoFisher Dionex Ultimate 3000 RSLCnano system. All MS/MS analyses were performed in data independent acquisition mode.

The raw MS/MS data was processed using MaxQuant (version v2.6.3.0) to generate iBAQ values for select proteins. The amino acid sequences for the following target proteins were provided for the analysis: VP1, GFPnb, ALFAnb, GFP, GFP-NLS, and GFP-NLS-ALFA. A list of contaminant proteins derived from *Homo sapiens* found in samples purified from mammalian cells and common MS contaminants were also provided. Oxidation and acetyl protein N-term were included as variable modifications and carbamidomethyl as a fixed modification. A custom R script was used to extract the iBAQ values for each sample corresponding to the target proteins listed above. The GFP:capsid ratio was calculated using the equation below:

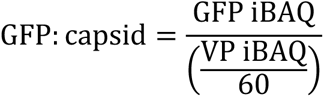

### Pulldown assay

All steps of the pulldown assay were performed with ice cold buffer and incubations were done at 4 °C unless otherwise stated. First, 20 µL of GFP Selector bead slurry (NanoTag Biotechnologies) were washed three times with 1 mL cold PBS to equilibrate the beads. Next, the beads were resuspended in 490 µl cold PBS and 10 µL recombinant GFP protein with a C-terminal ALFA tag (GFP-ALFA, NanoTag Biotechnologies) was added. To allow for binding of the GFP-ALFA protein to the beads, samples were incubated for 1 hour in an end-over-end rotator. Next, samples were washed three times with 1 mL PBS and afterwards with 1 mL TBS by pipetting up and down a couple of times to eliminate unbound GFP-ALFA protein. The beads were then resuspended in 490 µL PBS and 10 µL of iodixanol gradient purified AAV were added. Samples were incubated for another hour in an end-over-end rotator. Samples were washed again three times with 1 mL PBS followed by 1 mL TBS before they were resuspended in 50 µL PBS. To extract the ssDNA payload from the AAV particles bound to the beads, 5.5 µL Proteinase K buffer and 1 µL Proteinase K were added and samples incubated for 20 min at 50 °C. Proteinase K was then heat inactivated by an incubation at 95 °C for 10 min before the ssDNA was extracted using the DNA Clean & Concentrator-5 Kit. AAV samples from before and after the pulldown were then quantified using qPCR as described above. Percentage of bound AAV were calculated and normalized to the wtAAV-DJ control.

### APEX activity assay

To test for packaged peroxidase activity, the AMPLEX Red Hydrogen Peroxide Assay kit was used (Invitrogen). AMPLEX reaction mix was prepared according to manufacturer’s instructions and 50 μL of this was added to each well of a 96 well glass bottom plate (Greiner Bio-One). 10 μL of iodixanol purified virus was then added to each well and the reaction was incubated in the dark at room temperature for 30 minutes. Fluorescence was detected using a Synergy HTX multi-well plate reader with excitation/emission at 540nm/600nm respectively. 1x Reaction Buffer and WT-AAV-DJ were used as negative controls, while dilutions of Horseradish Peroxidase (HRP) from the kit was used as a positive control. Two biological replicates and at least two technical replicates were performed for each sample.

### Statistics

For the live cell imaging data, the normality of the dataset was assessed using the Shapiro-Wilk test which indicated a non-normal distribution. The non-parametric Kruskal-Willis test was used to test for statistical difference between the indicated variants followed by a post hoc Dunn’s test with Benjamini-Hochberg correction for comparison to the indicated control. For the APEX2 activity data, a Dunnett’s test was used to test for statistical difference between the indicated variants for comparison to the indicated control. *p*-values <0.05 were considered statistically significant (\**p* < 0.05; \*\**p* < 0.01; \*\*\**p* < 0.001). All *p*-values are listed in **Table S2** and **Table S3**. All statistical analyses were performed in R (version 4.3.3).

## DATA AVAILABILITY

All plasmids are available upon request.

## AUTHOR CONTRIBUTIONS

MDH, RS, AE, and DS designed the study. MDH, RS, and AE conducted the experiments with assistance from YH. MDH, RS, AE, and DS analyzed the data and authored the manuscript. All authors have given approval to the final version of the manuscript.

## ACKNOWLEDGEMENTS

This work was also supported by the resources and staff at the University of Minnesota University Imaging Centers (UIC). The UIC RRID is SCR_020997. We thank the Center for Metabolomics and Proteomics at the University of Minnesota for providing services related to generation of quantitative mass spectrometry data. pAAV-CAG-GFP (Addgene plasmid #37825) and pAAV-CAG-tdTomato (codon diversified; Addgene plasmid # 59462) were a gift from Edward Boyden, pATT-Dest was a gift from David Savage (Addgene plasmid #79770), APEX2-NLS was a gift from Alice Ting (Addgene plasmid #124167), and pdCas9-GFP was a gift from Karsten Rippe (Addgene plasmid #181906).

## FUNDING

The project described was supported by a Career Development Award from the American Society of Gene & Cell Therapy. The content is solely the responsibility of the authors and does not necessarily represent the official views of the American Society of Gene & Cell Therapy. This work was supported by the National Institutes of Health (R01GM141152 to DS & AL) and (P30DA048742-01A1 to DS).

## CONFLICT OF INTEREST

MDH and DS have filed a patent application related to this study (PCT/US2024/025273). All other authors state no conflict of interest.

