## Supplementary Materials for "Protein Carrier AAV"

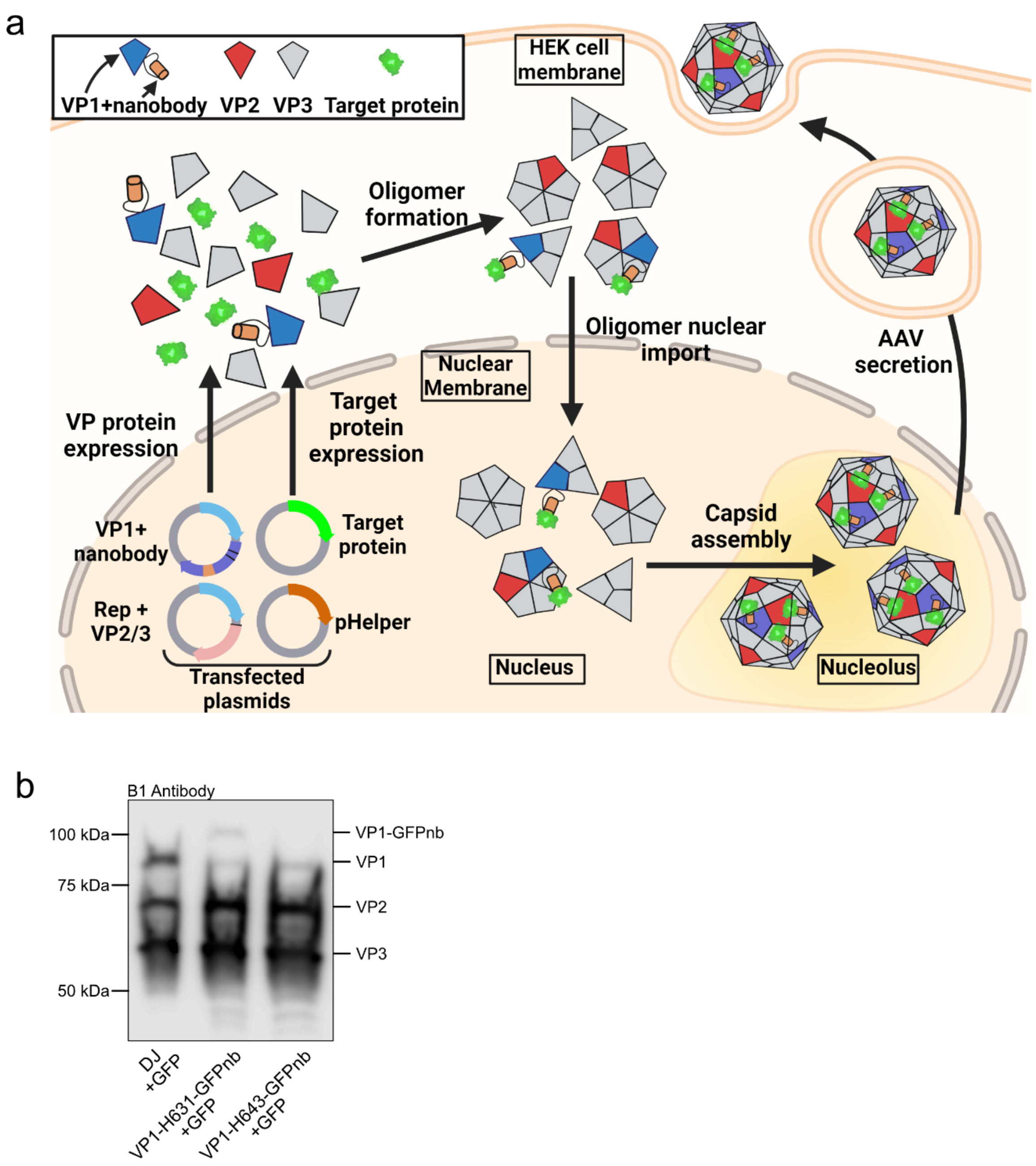


**Figure S1: Proposed packaging mechanism of a target protein by engineered AAV capsids and characterization of VP1 GFP-nanobody (GFPnb) subunit incorporation. (a)** Cells are simultaneously transfected with plasmids encoding for a target protein, a VP protein containing a nanobody insertion, complementary VP proteins for capsid assembly, and pHelper. The VP containing the nanobody insertion associates with the target protein during oligomer assembly in the cytoplasm or capsid assembly in the nucleus resulting in AAV capsids packaging proteins. **(b)** Western blot analysis demonstrating that VP1 subunits with a GFPnb inserted at H631 or H643 are incorporated into assembled capsids. Samples were run on a 7.5% SDS-PAGE gel to better distinguish between the VP subunits.

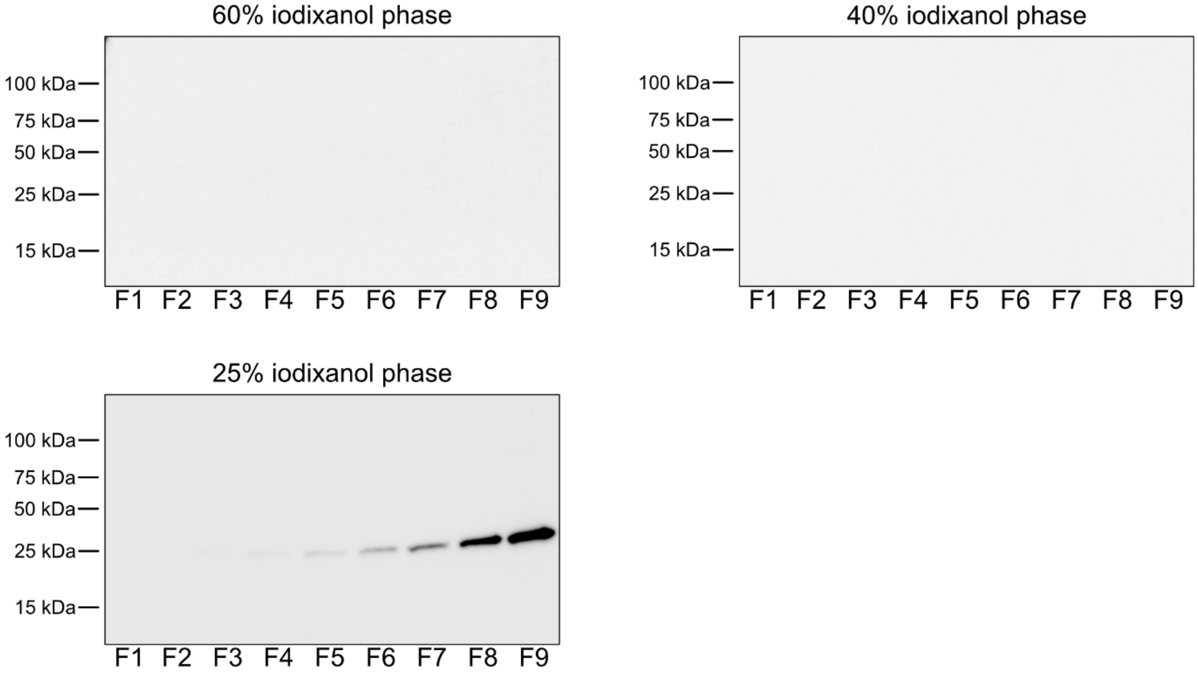


**Figure S2: GFP ends up in the upper 25 % iodixanol gradient phase post ultracentrifugation.** Western blot with GFP staining from nine fractions (F1-F9) of each iodixanol phase (60 %, 40 %, and 25 %).

a

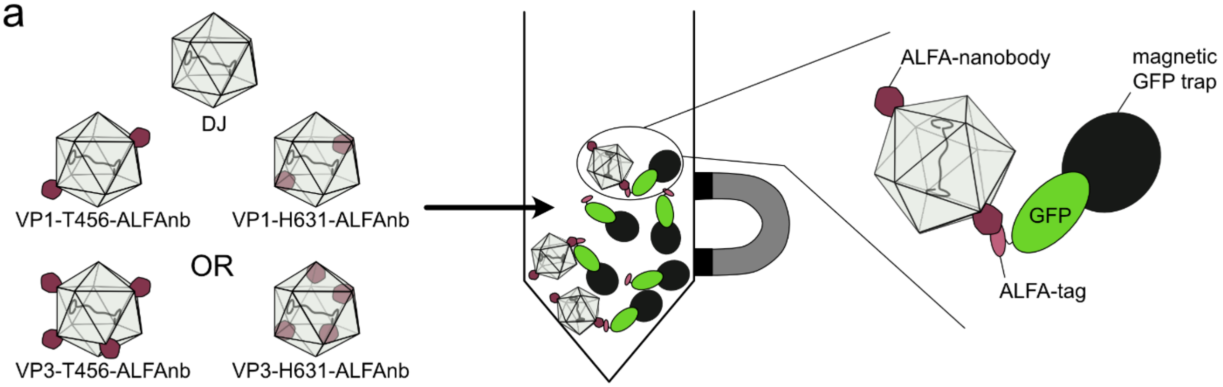


b

| sample | % pulled down |
|---|---|
| DJ | 0.00 |
| VP1-H631-ALFAnb | 3.05 |
| VP1-T456-ALFAnb | 10.66 |
| VP3-H631-ALFAnb | 0.00 |
| VP3-T456-ALFAnb | 16.57 |

**Figure S3: Pulldown of capsids with ALFA-nanobody (ALFAnb) insertions on the capsid's surface. (a)** Schematic of pulldown whereas the magnetic GFP trap first captures a GFP with a C-terminal ALFA tag, which in a second step binds to the ALFA-nanobody incorporated into the outside of the AAV capsid. **(b)** Table listing the percentages of pulled down AAV particles normalized to the DJ control. Data are means of two replicates.

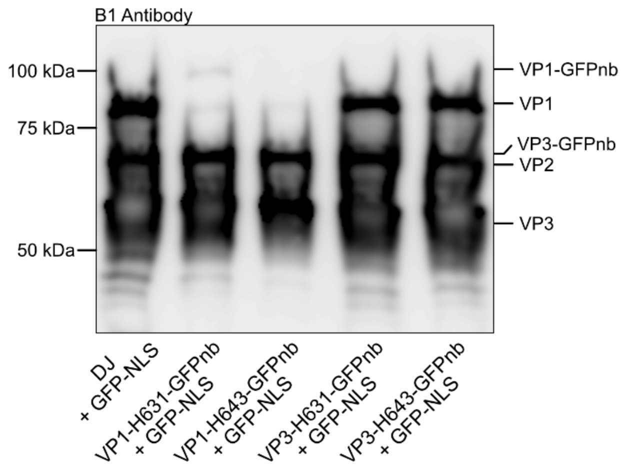


**Figure S4: Western blot showing the degree of incorporation of VP1 or VP3 with a GFPnb inserted at H631 or H643 into assembled capsids.** The samples were run on a 7.5% gel to better visualize the VP subunits, however, the VP3 GFPnb band does overlap with that of VP2 due to the similar molecular weights of the two proteins (VP3 GFPnb = 70.4 kDa and VP2 = 66.6 kDa).

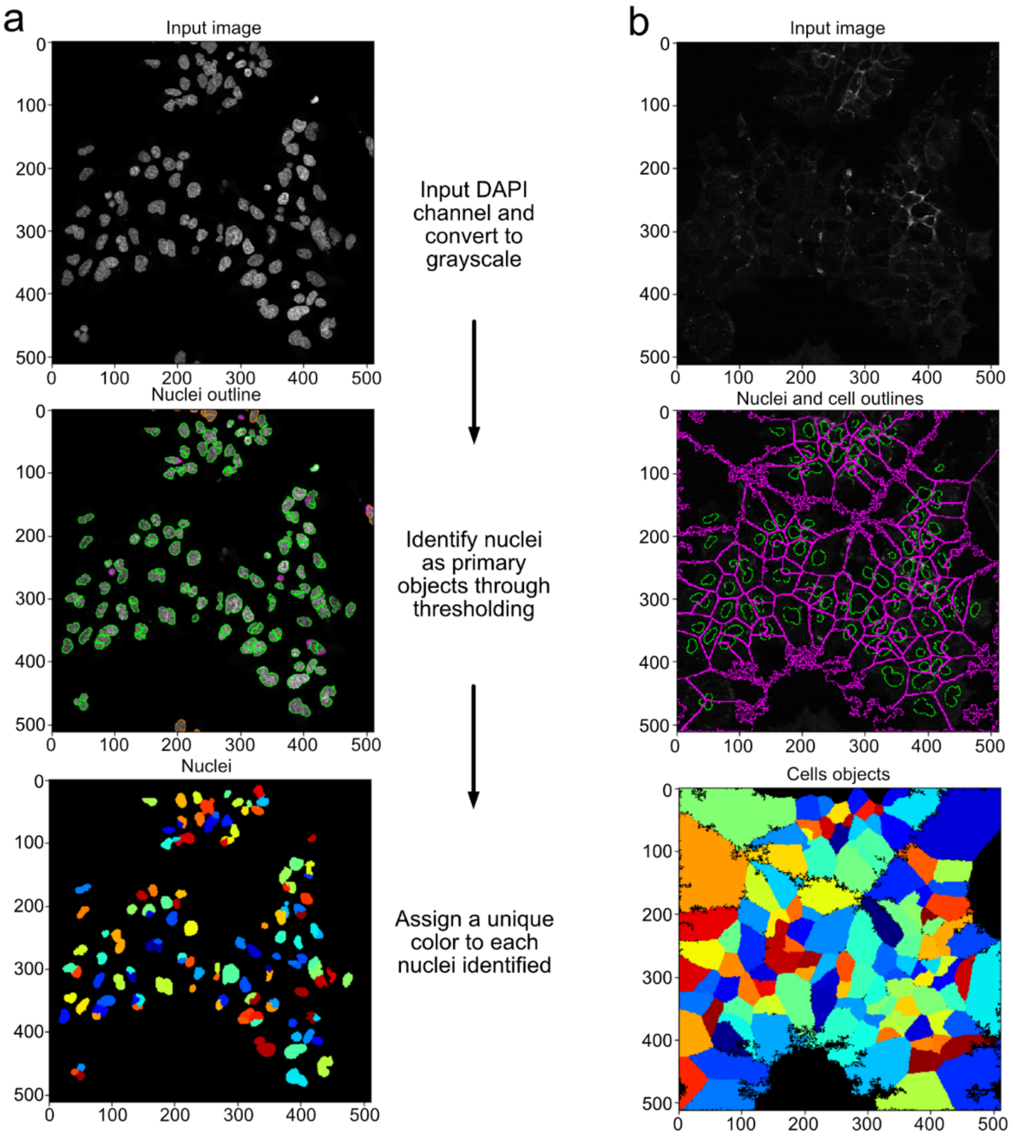


**Figure S5: Automated imaging pipeline representative images. (a)** Nuclei are first identified as primary objects using the DAPI channel. **(b)** The identified nuclei are subsequently used to identify cells as secondary objects using the WGA Alexa Fluor 555 channel.

a

GFP nanobody with linkers

**SGGGGQVQLVESGGALVQPGGSLRLSCAASGFPVNRYSMRWYRQAPGKEREWV
AGMSSAGDRSSYEDSVKGRFTISRDDARNTVYLQMNSLKPEDTAVYYCNVNVGFE
YWGQGTQVTVSSKGGGGS**

ALFA nanobody with linkers

**SGGGGSGEVQLQESGGGLVQPGGSLRLSCTASGVTISALNAMAMGWYRQAPGE
RRVMVAAVSERGNAMYRESVQGRFTVTRDFTNKMVSLQMDNLKPEDTAVYYCHV
LEDRVDSFHDYWGQGTQVTVSSGGGGS**

b

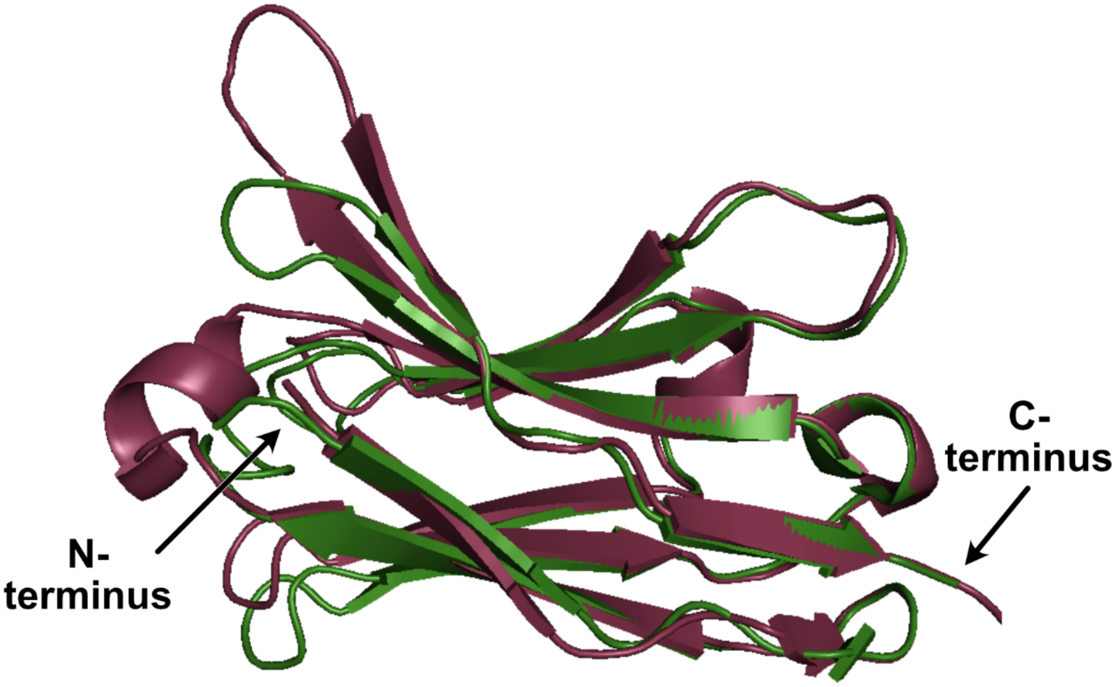


**Figure S6: GFP and ALFA nanobodies. (a)** Protein sequences of GFP (dark green) and ALFA (ruby) nanobodies with linkers (yellow). **(b)** Structure alignment of GFP and ALFA (RCSB PDB 3K1K) nanobodies (RCSB PDB 6I2G).

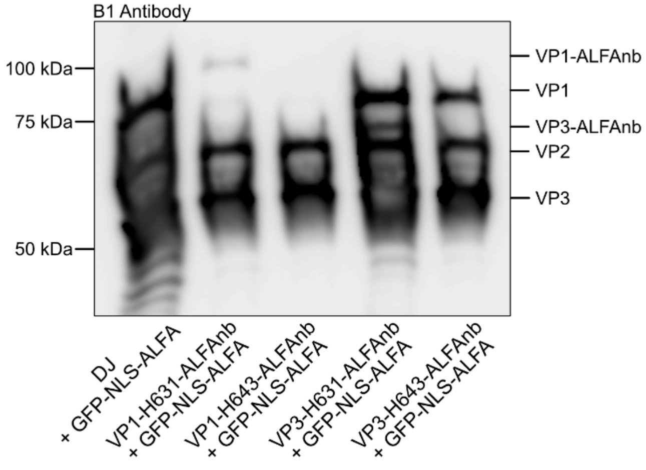


**Figure S7:** Western blot showing the degree of incorporation of VP subunits containing an ALFAnb insertion at either position H631 or H643. For both VP1 and VP3, insertion of the ALFAnb at position H631 enables increased incorporation into assembled capsids when compared to insertion at H643.

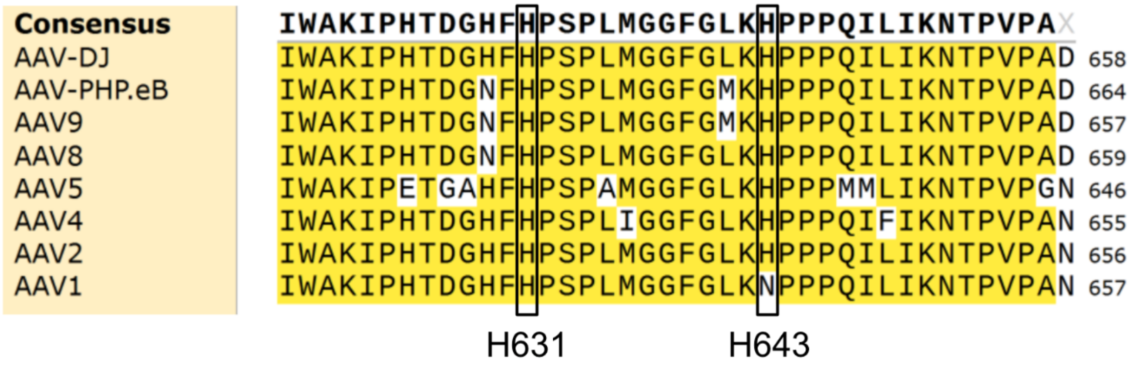


**Figure S8:** Sequence alignment of AAV serotypes AAV-DJ, AAV-PHP.eB, AAV9, AAV8, AAV5, AAV4, AAV2, and AAV1 with the two binding domain insertion positions highlighted H631 and H643 (black rectangles). Residue H631 is conserved across all serotypes shown, while residue H643 is conserved in all serotypes except AAV1.

**GFP-NLS-ALFA-tag**

MVSKGEELFTGVVPILVELDGDVNGHKFSVSGEGEGDATYGKLTLKFICTTGKLPVPWPTLVT
TLTYGVQCFSRYPDHMKQHDFFKSAMPEGYVQERTIFFKDDGNYKTRAEVKFEGDTLVNRIE
LKGIDFKEDGNILGHKLEYNYNSHNVYIMADKQKNGIKVNFKIRHNIEDGSVQLADHYQQNTPI
GDGPVLLPDNHYLSTQSALSKDPNEKRDHMVLLEFVTAAGITLGMDELYKSGGGGPAAKRV
KLDGGGGSPSRLEEELRRRLTE

**spCas9-GFP**

PKKKRKVGRVCRISSLRYRGPGIATMDKKYSIGLDIGTNSVGWAVITDEYKVPSKKFKVLGNT
DRHSIKKNLIGALLFDSGETAEATRLKRTARRRYTRRKNRICYLQEIFSNEMAKVDDSFFHRLE
ESFLVEEDKKHERHPIFGNIVDEVAYHEKYPTIYHLRKKLVDSTDKADLRLIYLALAHMIKFRG
HFLIEGDLNPDNSDVDKLFIQLVQTYNQLFEENPINASGVDAKAILSARLSKSRRLENLIAQLP
GEKKNGLFGNLIALSLGLTPNFKSNFDLAEDAKLQLSKDTYDDDLDNLLAQIGDQYADLFLAA
KNLSDAILLSDILRVNTEITKAPLSASMIKRYDEHHQDLTLLKALVRQQLPEKYKEIFFDQSKN
GYAGYIDGGASQEEFYKFIKPILEKMDGTEELLVKLNREDLLRKQRTFDNGSIPHQIHLGELHA
ILRRQEDFYPFLKDNREKIEKILTFRIPYYVGPLARGNSRFAWMTRKSEETITPWNFEEVVDKG
ASAQSFIERMTNFDKNLPNEKVLPKHSLLYEYFTVYNELTKVKYVTEGMRKPAFLSGEQKKAI
VDLLFKTNRKVTVKQLKEDYFKKIECFDSVEISGVEDRFNASLGTYHDLLKIIKDKDFLDNEEN
EDILEDIVLTLTLFEDREMIEERLKTYAHLFDDKVMKQLKRRRYTGWGRLSRKLINGIRDKQSG
KTILDFLKSDGFANRNFMQLIHDDSLTFKEDIQKAQVSGQGDSLHEHIANLAGSPAIKKGILQT
VKVVDELVKVMGRHKPENIVIEMARENQTTQKGQKNSRERMKRIEEGIKELGSQILKEHPVEN
TQLQNEKLYLYYLQNGRDMYVDQELDINRLSDYDVDHIVPQSFLKDDSIDNKVLTRSDKNRG
KSDNVPSEEVVKKMKNYWRQLLNAKLITQRKFDNLTKAERGGLSELDKAGFIKRQLVETRQI
TKHVAQILDSRMNTKYDENDKLIREVKVITLKSKLVSDFRKDFQFYKVREINNYHHAHDAYLN
AVVGTALIKKYPKLESEFVYGDYKVYDVRKMIAKSEQEIGKATAKYFFYSNIMNFFKTEITLAN
GEIRKRPLIETNGETGEIVWDKGRDFATVRKVLSMPQVNIVKKTEVQTGGFSKESILPKRNSD
KLIARKKDWDPKKYGGFDSPTVAYSVLVVAKVEKGKSKKLKSVKELLGITIMERSSFEKNPID
FLEAKGYKEVKKDLIIKLPKYSLFELENGRKRMLASAGELQKGNELALPSKYVNFLYLASHYE
KLKGSPEDNEQKQLFVEQHKHYLDEIIEQISEFSKRVILADANLDKVLSAYNKHRDKPIREQAE
NIIHLFTLTNLGAPAAFKYFDTTIDRKRYTSTKEVLDATLIHQSITGLYETRIDLSQLGGDAYPYD
VPDYAPVPKKKRKVPVATRILQSTVPRARDPPVATMVSKGEELFTGVVPILVELDGDVNGHKF
SVSGEGEGDATYGKLTLKFICTTGKLPVPWPTLVTTLTYGVQCFSRYPDHMKQHDFFKSAMP
EGYVQERTIFFKDDGNYKTRAEVKFEGDTLVNRIELKGIDFKEDGNILGHKLEYNYNSHNVYIM
ADKQKNGIKVNFKIRHNIEDGSVQLADHYQQNTPIGDGPVLLPDNHYLSTQSALSKDPNEKRD
HMVLLEFVTAAGITLGMDELYKTSVYNVQSGRDSRS

**Cre-ALFA**

MGPSRLEEELRRRLTEGGGGSPKKKRKVSNLLTVHQNLPALPVDATSDEVRKNLMDMFRDR
QAFSEHTWKMLLSVCRSWAAWCKLNNRKWFPAEPEDVRDYLLYLQARGLAVKTIQQHLGQ
LNMLHRRSGLPRPSDSNAVSLVMRRIRKENVDAGERAKQALAFERTDFDQVRSLMENSDRC
QDIRNLAFLGIAYNTLLRIAEIARIRVKDISRTDGGRMLIHIGRTKTLVSTAGVEKALSLGVTKLV
ERWISVSGVADDPNNYLFCRVRKNGVAAPSATSQLSTRALEGIFEATHRLIYGAKDDSGQRY
LAWSGHSARVGAARDMARAGVSIPEIMQAGGWTNVNIVMNYIRNLDSETGAMVRLLEDGD

**APEX2-NES**

MVRGSGKPIPNPLLGLDSTGKSYPTVSADYQDAVEKAKKKLRGFIAEKRCAPLMLRLAFHSA
GTFDKGTKTGGPFGTIKHPAELAHSANNGLDIAVRLLEPLKAEFPILSYADFYQLAGVVAVEV
TGGPKVPFHPGREDKPEPPPEGRLPDPTKGSDHLRDVFGKAMGLTDQDIVALSGGHTIGAA
HKERSGFEGPWTSNPLIFDNSYFTELLSGEKEGLLQLPSDKALLSDPVFRPLVDKYAADEDA
FFADYAEAHQKLSELGFADALQLPPLERLTLD

**APEX2-cMyc-NLS**
MVRGSGKPIPNPLLGLDSTGKSYPTVSADYQDAVEKAKKKLRGFIAEKRCAPLMLRLAFHSA
GTFDKGTKTGGPFGTIKHPAELAHSANNGLDIAVRLLEPLKAEFPILSYADFYQLAGVVAVEV
TGGPKVPFHPGREDKPEPPPEGRLPDPTKGSDHLRDVFGKAMGLTDQDIVALSGGHTIGAA
HKERSGFEGPWTSNPLIFDNSYFTELLSGEKEGLLQLPSDKALLSDPVFRPLVDKYAADEDA
FFADYAEAHQKLSELGFADASGGGGPAAKRVKLD

**APEX2-ALFA**
MVRGSGKPIPNPLLGLDSTGKSYPTVSADYQDAVEKAKKKLRGFIAEKRCAPLMLRLAFHSA
GTFDKGTKTGGPFGTIKHPAELAHSANNGLDIAVRLLEPLKAEFPILSYADFYQLAGVVAVEV
TGGPKVPFHPGREDKPEPPPEGRLPDPTKGSDHLRDVFGKAMGLTDQDIVALSGGHTIGAA
HKERSGFEGPWTSNPLIFDNSYFTELLSGEKEGLLQLPSDKALLSDPVFRPLVDKYAADEDA
FFADYAEAHQKLSELGFADAGGGGSPSRLEEELRRRLTE

**APEX2-cMyc-NLS-ALFA**
MVRGSGKPIPNPLLGLDSTGKSYPTVSADYQDAVEKAKKKLRGFIAEKRCAPLMLRLAFHSA
GTFDKGTKTGGPFGTIKHPAELAHSANNGLDIAVRLLEPLKAEFPILSYADFYQLAGVVAVEV
TGGPKVPFHPGREDKPEPPPEGRLPDPTKGSDHLRDVFGKAMGLTDQDIVALSGGHTIGAA
HKERSGFEGPWTSNPLIFDNSYFTELLSGEKEGLLQLPSDKALLSDPVFRPLVDKYAADEDA
FFADYAEAHQKLSELGFADASGGGGPAAKRVKLDGGGGSPSRLEEELRRRLTE

**APEX2-SV40-NLS-ALFA**
MVRGSGKPIPNPLLGLDSTGKSYPTVSADYQDAVEKAKKKLRGFIAEKRCAPLMLRLAFHSA
GTFDKGTKTGGPFGTIKHPAELAHSANNGLDIAVRLLEPLKAEFPILSYADFYQLAGVVAVEV
TGGPKVPFHPGREDKPEPPPEGRLPDPTKGSDHLRDVFGKAMGLTDQDIVALSGGHTIGAA
HKERSGFEGPWTSNPLIFDNSYFTELLSGEKEGLLQLPSDKALLSDPVFRPLVDKYAADEDA
FFADYAEAHQKLSELGFADAEFSRADPKKKRKVDPKKKRKVDPKKKRKVGGGGSPSRLEE
ELRRRLTE

**Figure S9:** Amino acid sequences of select packaged proteins. GFP sequence in light green, spCas9 in dark blue, Cre in dark green, linkers in brown, NLS in light pink, and ALFA tag in dark pink.

**Hoffmann et al. - uncropped gel images**

**Figure 1c**

B1 blot

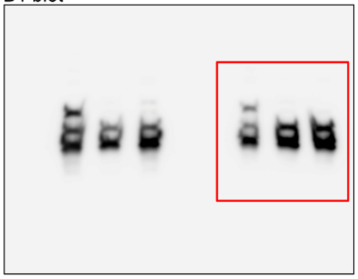

GFP blot

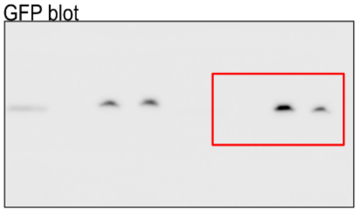

**Figure 2a**

B1 blot

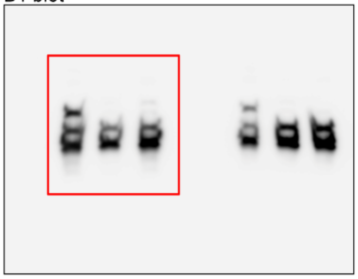

GFP blot

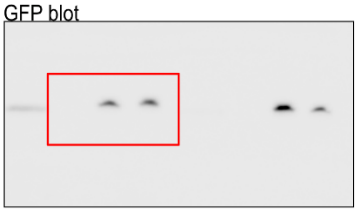

**Figure 2c**

B1 blot

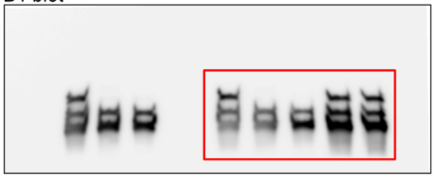

GFP blot

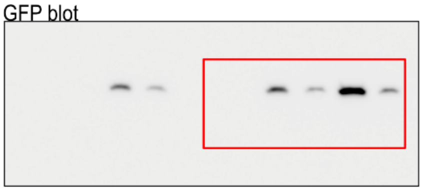

**Figure 4a**

B1 blot

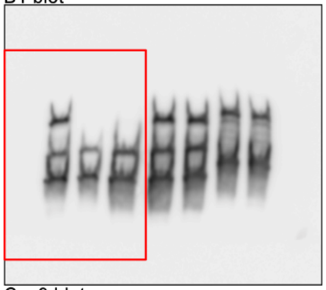

Cas9 blot

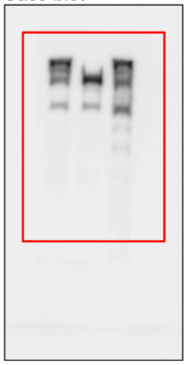

**Figure 4b**

B1 blot

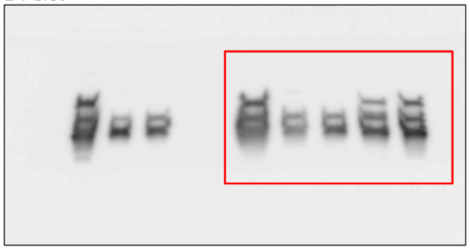

GFP blot

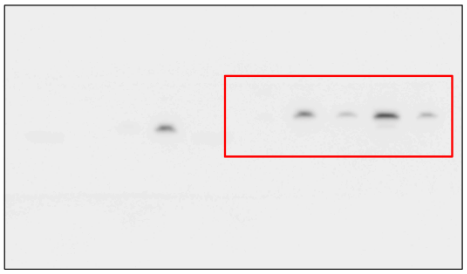

**Figure 4c**

B1 blot

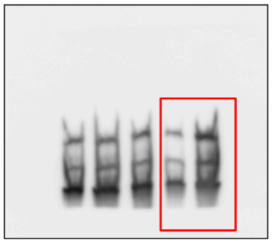

ALFA blot

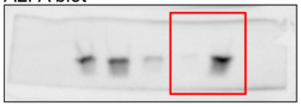

**Figure 4d**

B1 blot

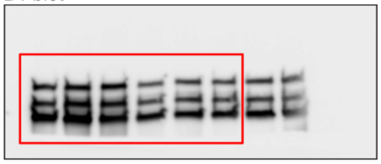

V5 blot

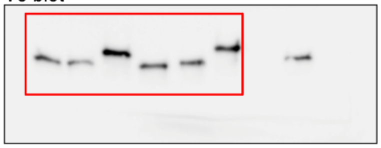

**Hoffmann et al. - uncropped gel images**

**Figure S1b**
B1 blot

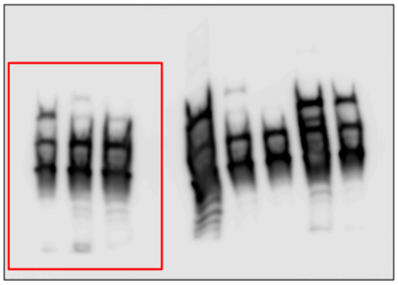

**Figure S2**
GFP blot (60%)

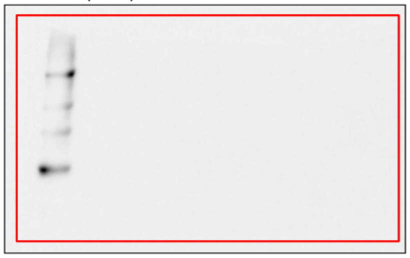

GFP blot (40%)

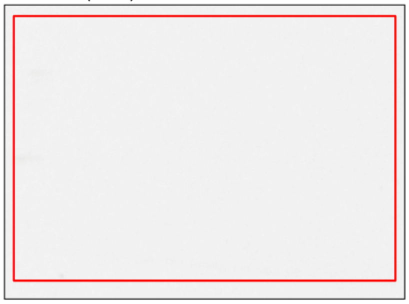

GFP blot (25%)

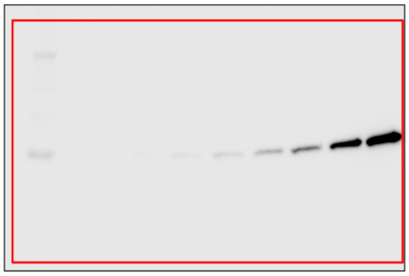

**Figure S4**
B1 blot

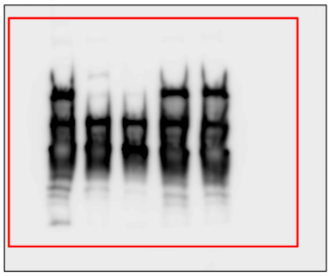

**Figure S7**
B1 blot

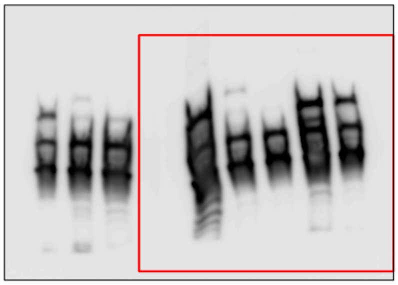

**Figure S10:** Uncropped Western blots for all figures.

**Table S1**
Plasmids used in this study.

| # | Use | Name | Expressed Genes | Origin |
|---|---|---|---|---|
| 1 | | GFP | GFP | Schmidt Lab MDH |
| 2 | | GFP-NLS | GFP-NLS | Schmidt Lab RS |
| 3 | | GFP-NLS-ALFA | GFP-NLS-ALFA | Schmidt Lab RS |
| 4 | | spCas9 | spCas9 | Schmidt Lab MDH |
| 5 | | spCas9-GFP | spCas9-GFP | Schmidt Lab MDH |
| 6 | | ALFA-Cre | ALFA-SV40-NLS-Cre | Schmidt Lab MDH |
| 7 | cargo plasmids | APEX2 | APEX2 | Addgene #124617 |
| 8 | | APEX2-NES | APEX2-NES | Schmidt Lab AE |
| 9 | | APEX2-cMyc-NLS | APEX2-cMyc-NLS | Schmidt Lab AE |
| 10 | | APEX2-ALFA | APEX2-ALFA | Schmidt Lab AE |
| 11 | | APEX2-cMyc-NLS-ALFA | APEX2-cMyc-NLS-ALFA | Schmidt Lab AE |
| 12 | | APEX2-SV40-NLS-ALFA | APEX2-SV40-NLS-ALFA | Schmidt Lab AE |
| 13 | | ITR-tdTomato | tdTomato | Addgene #59462 |
| 14 | AAV production complementation plasmids | pHelper | Adeno helper genes | CellBiolabs |
| 15 | | wt DJ | rep2 + capDJ | CellBiolabs |
| 16 | | DJ-M1K | rep2 + capDJ-VP2/VP3 | Schmidt Lab AZ |
| 17 | GFP-nanobody insertion | DJ-VP1-H631-GFPnb | rep2 + VP1-H631-GFPnb | Schmidt Lab MDH |
| 18 | | DJ-VP1-H643-GFPnb | rep2 + VP1-H643-GFPnb | Schmidt Lab MDH |
| 19 | | DJ-VP3-H631-GFPnb | rep2 + VP3-H631-GFPnb | Schmidt Lab MDH |
| 20 | | DJ-VP3-H643-GFPnb | rep2 + VP3-H643-GFPnb | Schmidt Lab MDH |
| 21 | ALFA-nanobody insertion | DJ-VP1-H631-ALFAnb | rep2 + VP1-H631-ALFAnb | Schmidt Lab MDH |
| 22 | | DJ-VP1-H643-ALFAnb | rep2 + VP1-H643-ALFAnb | Schmidt Lab MDH |
| 23 | | DJ-VP1-T456-ALFAnb | rep2 + VP1-T456-ALFAnb | Schmidt Lab MDH |
| 24 | | DJ-VP3-H631-ALFAnb | rep2 + VP3-H631-ALFAnb | Schmidt Lab MDH |
| 25 | | DJ-VP3-H643-ALFAnb | rep2 + VP3-H643-ALFAnb | Schmidt Lab MDH |
| 26 | | DJ-VP3-T456-ALFAnb | rep2 + VP3-T456-ALFAnb | Schmidt Lab MDH |

**Table S2**

p-values of post hoc Dunn's test with Benjamini-Hochberg correction from Figure 3b.

| treatment condition | p value | significance |
|---|---|---|
| GFP | 0.5261 | n.s. |
| empty wtAAV-DJ | 3.8434e-10 | *** |
| VP3-H631-GFPnb + GFP-NLS | <2e-16 | *** |

**Table S3**

p-values of Dunnett's test from Figure 4e.

| variant | p value | significance |
|---|---|---|
| DJ + APEX2 | 0.987 | n.s. |
| DJ + APEX2-NES | 0.990 | n.s. |
| VP3-H643-ALFAnb + APEX2-ALFA | 0.982 | n.s. |
| DJ + APEX2-cMyc | 0.000238 | *** |
| VP3-H643-ALFAnb + APEX2-cMyc-ALFA | 1.00 | n.s |
| VP3-H643-ALFAnb + APEX2-SV40-ALFA | 0.787 | n.s |
